# A-to-I RNA editing uncovers hidden signals of adaptive genome evolution in animals

**DOI:** 10.1101/228734

**Authors:** Niko Popitsch, Christian D. Huber, Ilana Buchumenski, Eli Eisenberg, Michael Jantsch, Arndt von Haeseler, Miguel Gallach

## Abstract

In animals, the most common type of RNA editing is the deamination of adenosines (A) into inosines (I). Because inosines base-pair with cytosines (C), they are interpreted as guanosines (G) by the cellular machinery and genomically encoded G alleles at edited sites mimic the function of edited RNAs. The contribution of this hardwiring effect on genome evolution remains obscure. We looked for population genomics signatures of adaptive evolution associated with A-to-I RNA edited sites in humans and *Drosophila melanogaster*. We found that single nucleotide polymorphisms at edited sites occur 3 (humans) to 15 times (*Drosophila*) more often than at unedited sites, the nucleotide G is virtually the unique alternative allele at edited sites and G alleles segregate at higher frequency at edited sites than at unedited sites. Our study reveals that coding synonymous and nonsynonymous as well as silent and intergenic A-to-I RNA editing sites are likely adaptive in the distantly related human and *Drosophila* lineages.

## Introduction

Through a single nucleotide modification, A-to-I RNA editing may impact the stability of the corresponding RNA molecule, recode the original protein sequence, and eventually modulate its biological function. The role of RNA editing in animal evolution is not well understood. A widely accepted hypothesis suggests that A-to-I RNA editing at nonsynonymous sites would entail a selective advantage over a genomic G nucleotide, as it increases the transcriptome diversity without affecting the genomically encoded A phenotype in tissues where editing does not occur[1-3]. This hypothesis predicts that edited A nucleotide sites will be rarely substituted by G nucleotides compared to unedited A sites (hypothesis H1, Table 1). Contrary to this prediction, it was shown that A-to-G nucleotide substitutions between species are more frequent at edited sites than at unedited sites[4,5]. An alternative hypothesis (hypothesis H2, Table1) suggests that nonsynonymous A-to-G nucleotide substitutions between species are more tolerated (i.e., less deleterious) at edited sites than at unedited sites[4], explaining the difference in A-to-G substitution rates. Finally, a third hypothesis (hypothesis H3, Table1) proposes that G nucleotide sites are the ancestral state of currently edited A sites, and that A-to-I RNA editing is a compensation mechanism to reverse the harmful A phenotype caused by G-to-A mutations[5-7]. However, the fact that the editing level is far below 100% (for instance, in *D. melanogaster* the average editing level is 23%[8]) suggests that A-to-I RNA editing would rarely overcome the deleterious effects of the G-to-A mutations. In any case, each hypothesis predicts different evolutionary outcomes for the non-synonymous edited sites compared to unedited sites (Table 1).

**Table 1.** Hypotheses suggested for the evolution of A-to-I RNA editing target sites

|  | Hypothesis |  |  |  |
| --- | --- | --- | --- | --- |
|  | H1: Transcriptome diversity is beneficial (1-3) | H2: G is slightly deleterious (4) | H3: Compensatory hypothesis (5-7) | H4: Adaptive hypothesis (current study) |
| **Features** |  |  |  |  |
| Ancestral state | A | A | G | A |
| Adaptive value of editing | Editing is adaptive because provides diversity to transcript population. | Editing is very deleterious and currently detected edited sites are generally slightly deleterious. | Editing is adaptive as it reverses the harmful effect of G-to-A mutations. | Editing is adaptive because A-to-I replacements are beneficial at these nucleotide sites. |
| Relative fitness (S) of the derived allele | $S_A > S_G \geq S_{C,T}$ | $S_A \geq S_G \geq S_{C,T}$ | $S_G \geq S_A > S_{C,T}$ | $S_G > S_A \gg S_{C,T}$ |
| **Population genetics predictions compared to unedited sites** |  |  |  |  |
| Overall polymorphic rate | Polymorphism at edited sites should be **reduced** as A-to-G, A-to-C and A-to-T mutations are slightly deleterious. | Polymorphism at edited sites should be **slightly increased** as A-to-G mutations are slightly more tolerated than at unedited sites. | Polymorphism at edited sites should be **similar or slightly increased** as editing somehow reduces the deleterious effect of G-to-A mutations. | Polymorphism at edited sites should be **increased** as A-to-G mutations are largely adaptive. |
| Polymorphism type | **A,G should be slightly more frequent** than A,C and A,T polymorphisms at edited sites. | **A,G should be slightly more frequent** than A,C and A,T polymorphisms at edited sites. | **A,G should be slightly more frequent** than A,C and A,T polymorphisms at edited sites. | **A,C and A,T polymorphism should be rarely found.** |
| Polymorphic rate at coding regions | **Similar or reduced** at both edited and unedited sites due to potential deleterious effects at non-synonymous sites. | **Similar or reduced** at both edited and unedited sites due to potential deleterious effects at non-synonymous sites. | **Similar or reduced** at both edited and unedited sites due to potential deleterious effects at non-synonymous sites. | **Increased** at edited sites as the G allele mimics the protein variant obtained through editing. |
| Synonymous polymorphic rate | **Similar** at both edited and unedited sites. | **Similar** at both edited and unedited sites. | **Similar** at both edited and unedited sites. | **Increased** at edited sites. |
| Frequency spectrum of the derived allele | **Derived G** allele should segregate at **similar or lower** frequency (i.e., purifying selection or neutral at most). | **Derived G** allele should segregate at **similar** frequency (i.e., neutral or nearly neutral). | **Derived G** allele should segregate at **similar** frequency (i.e., neutral or nearly neutral). | **Derived G** allele should segregate at **higher** frequency. |
| Nucleotide diversity around edited sites | **Similar** | **Similar** | **Similar** | **Reduced** |

To our knowledge, most studies have applied a phylogenetic approach to detect footprints of adaptive evolution of A-to-I RNA editing at coding regions[9-12]. Here, we employ a population genomics approach to search for signatures of selection in both coding and noncoding regions of the genome. To this end, we integrated the *D. melanogaster* and human editomes into population genomics data and investigated the population genetic patterns of the A-to-I RNA editing sites. Our study contradicts several predictions from previously suggested hypotheses and suggests a new adaptive role of A-to-I RNA editing in *Drosophila* and humans.

## Results

### Polymorphism patterns suggest adaptive editing in *Drosophila*

We analyzed *D. melanogaster* genome data from the *Drosophila* Genetics Reference Panel 2 (DGRP2)[13], consisting of 205 sequenced inbred lines derived from Raleigh (NC), U. S. A., and two additional wild populations collected in Florida (FL) and Maine (ME), U. S. A., consisting of 39 and 86 pool-sequenced inbred lines, respectively[14]. We investigated genome-wide nucleotide polymorphisms across more than 171 million nucleotide sites, 3,581 of them corresponding to known edited sites occurring in 1,074 genes[8]. We found that 15% (FL and ME) to 21% (DGRP2) of the edited sites are polymorphic, in sharp contrast to the 1% to 2% found among unedited sites (Table 2). This result does not support hypothesis H1 (Table 1), which predicts reduced polymorphisms at edited sites, but may be compatible with the hypotheses H2 and H3 (Table 1) which predict similar or slightly increased polymorphism at edited sites. Thus, according to the original study from where hypothesis H2 is derived[4], A-to-G nonsynonymous substitutions at edited sites are twice as frequent compared to nonsynonymous unedited sites (6.92% / 2.98% = 2.32). Although this study[4] compares humans and mice (not *Drosophila*), the 2.32-fold difference is far below the 10-(DGRP2) to 15-fold (FL and ME) increase in polymorphic rate at edited sites. We did not find a clear quantitative prediction for hypothesis H3[5-7]. Remarkably, we found that the G nucleotide is the alternative allele in at least 98% of the polymorphic edited sites (including both silent and non-synonymous ones), but only in ∼47% of the unedited polymorphic sites (Table 2). The percentage of each polymorphism type at unedited sites fits the transition (A-to-G) and transversion mutation (A-to-C and A-to-T) frequencies in Drosophila[15]. This result seems incompatible with hypotheses H1-H3 as all polymorphism types should be found, at least at silent edited sites (Table 1).

**Table 2.** Number of single nucleotide polymorphism sites and polymorphism types among edited and unedited sites in *Drosophila* populations and human.

|  | DGRP2 |  | Florida |  | Maine |  | Human - genic<sup>c</sup> |  | Human - intergenic<sup>c</sup> |  |
| --- | --- | --- | --- | --- | --- | --- | --- | --- | --- | --- |
|  | edited | unedited<sup>a</sup> | edited | unedited | edited | unedited | edited | unedited | edited | unedited |
| **Polymorphic** | **755 (21%)** | 3,951,070 (2%) | **543 (15%)** | 1,367,160 (1%) | **507 (14%)** | 1,235,454 (1%) | **231 (19%)** | 176,080 (6%) | **110 (18%)** | 196,030 (6%) |
| **Not polymorphic** | 2,826 (79%) | 171,048,930 (98%) | 3,038 (85%) | 118,920,513 (99%) | 3,074 (86%) | 119,052,219 (99%) | 977 (81%) | 2,811,804 (94%) | 520 (82%) | 3,017,246 (94%) |
| **Polymorphism<sup>a</sup> A,G** | **740 (98%)** | 817,333 (45%) | **536 (99%)** | 337,098 (48%) | **502 (99%)** | 309,347 (49%) | **225 (97%)** | 102,842 (58%) | **105 (96%)** | 112,936 (58%) |
| **A,C** | 3 (0%) | 355,952 (20%) | 1 (0%) | 142,183 (21%) | 0 (0%) | 131,624 (20%) | 4 (2%) | 35,491 (21%) | 3 (3%) | 38,772 (20%) |
| **A,T** | 12 (2%) | 649,599 (35%) | 6 (1%) | 217,230 (31%) | 5 (1%) | 195,528 (31%) | 2 (1%) | 37,747 (21%) | 2 (1%) | 42,971 (22%) |

These observations hold two important implications: 1) because C and T alleles are virtually absent at edited sites, A-to-I RNA editing is functionally constrained and likely adaptive relative to C and T, and 2) unless the A-to-G mutation rate is much higher at edited sites than at unedited sites due to an unknown molecular mechanism, the 10 to 15-fold increase in nucleotide polymorphism indicate that the G allele is likely adaptive at edited sites (hypothesis H4, Table1). We thus looked for additional evidence supporting the adaptive hypothesis.

### Derived G alleles at edited sites are likely adaptive in *Drosophila*

Among the 3,581 edited sites in *Drosophila*, 1,015 are protein coding nucleotides. Because of the potential deleterious effects caused by mutations in coding regions, nucleotide polymorphisms in such regions are expected to be similar or even lower than in noncoding regions[16]. This is what we see for unedited sites, where nucleotide polymorphisms remain at 2% (DGRP2) or even decreases from 1% to 0.5% (FL and ME; S1 Table). In contrast, nucleotide polymorphism at edited sites increases, on average, from 17% to 25% if we only consider coding regions. In other words, edited sites show a 16- to 44-times higher polymorphic rate than unedited sites at coding regions (S1 Table). This observation is not predicted by the hypotheses H1-H3 (Table 1) and prompted us to further investigate the relative contribution of nonsynonymous and synonymous replacements to nucleotide polymorphism at edited and unedited sites.

To understand the A,G polymorphism on a genome wide scale, we scanned the reference genome for coding A sites where a G mutation would result in a synonymous change. We found *S* = 777,461 A sites in the reference genome that would result in synonymous changes if replaced by G, 84,246 of which are actual synonymous A,G polymorphisms in the DGRP2 population, thus leading to a genomic rate of synonymous A,G polymorphisms 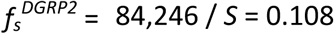. Similarly, we computed for edited sites the rate of synonymous A,G polymorphisms (251) per potentially synonymous A,G site (*S^edited^* = 370) as 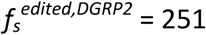 / *S^edited^* = 0.678. For the FL and ME populations we computed 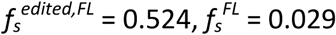 and 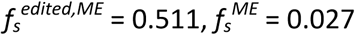, respectively. Therefore, the rate of synonymous A,G polymorphisms for edited sites is 6 to 19 times higher than for unedited sites in *Drosophila*. This result is rather inconsistent with hypotheses H1-H3 (Table 1) that predict similar rates of synonymous polymorphism at edited and unedited sites. Remarkably, for nonsynonymous sites, the differences between rates are even more pronounced: 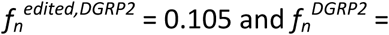 0. 007, which implies a 15-fold increased rate for edited nonsynonymous sites in DGRP2, while for the ME and FL populations the rate increase is 45-fold and 51-fold, respectively (Table 3).

**Table 3.**
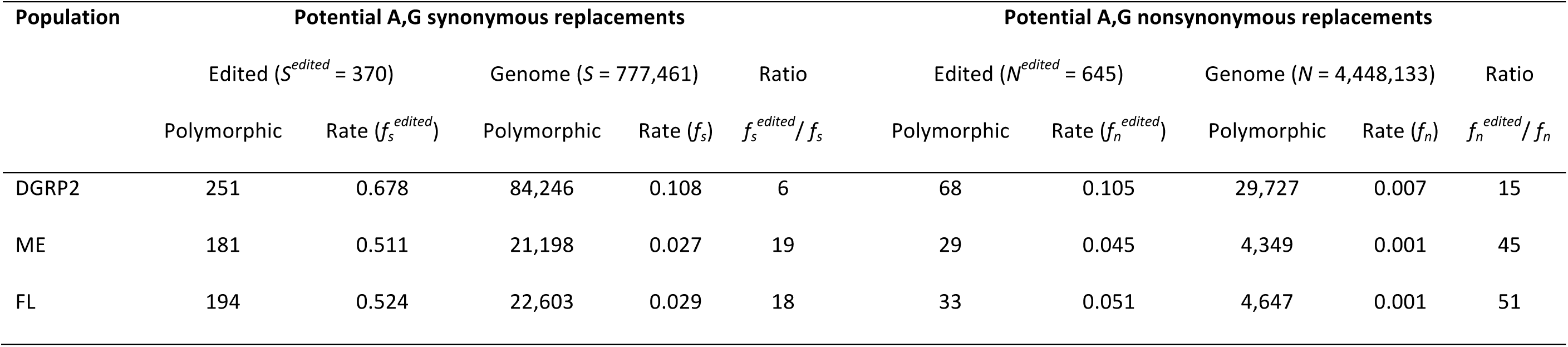
Potential A,G synonymous and nonsynonymous replacements in *Drosophila* populations.

A common way to determine the evolutionary force driving coding sequence evolution is the ratio of the number of nonsynonymous substitutions per nonsynonymous site (*d_N_*) to the number of synonymous substitutions per synonymous site (*d_S_*). The estimates of *f_s_* and *f_n_* fall within the distribution of *d_S_* (0.030 - 0.128; 5^th^ and 95^th^ percentiles, respectively) and *d_N_* (0.000 - 0.022; minimum and 95^th^ percentile, respectively) estimations for *D. melanogaster* genes[17]. We therefore applied the same reasoning behind the *d_N_* / *d_S_* ratio[18] to our *f_s_* and *f_n_* estimations. This is: if selection does not act on synonymous sites, then 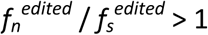 may be considered as an evidence of positive selection on nonsynonymous edited sites. However, the large polymorphism rate that we observe for edited sites and the fact that 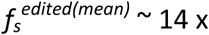 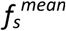 indicates that edited synonymous sites are not neutral but likely adaptive due to the pervasive roles of RNA editing in the posttranscriptional regulation of gene expression[19,20]. We therefore used 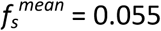 as the neutral rate for synonymous A,G polymorphisms in the genome, and obtained 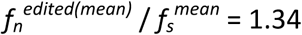 (*p* = 0.012, one-sided Binomial test for the null hypothesis 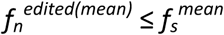). We conclude that the alleles encoding the same protein variant that is obtained through A-to-I RNA editing are likely adaptive.

According to population genetics theory, if the G alleles at polymorphic edited sites were adaptive, they would segregate at higher frequencies than G alleles at unedited sites originated at the same time[21]. This effect should be detectable by comparing the allele frequency spectrum for edited and unedited A,G polymorphisms. We used *D. simulans* population genomics data[16] to infer the ancestral state (i.e., polarize) of the polymorphic A-sites across the genome in the DGRP2 population and to be confident that the derived G alleles at edited and unedited sites are of similar age. We detected 462,498 A-to-G polymorphisms across the genome where the (derived) G allele most likely originated in *D. melanogaster’s* lineage, 303 of them occurring at edited sites (S2 Table). Fig 1a displays the allele frequency spectrum of the derived G alleles at edited and unedited A-to-G polymorphic sites. Remarkably, the frequency spectrum for the derived G alleles at edited sites is shifted to the right and quite distinct from that of unedited sites and from the expected allele frequency spectrum under neutral evolution, indicating that a significant fraction of A-to-G mutations at edited sites is likely adaptive. Our analysis in FL and ME populations supports this observation (S1 and S2 Figs). Because 266 (i.e., 88%) of the 303 polarized polymorphisms correspond to non-coding edited sites, the allele frequency spectrum analysis reveals a likely functional role of noncoding edited sites and endorses the use of 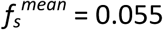 as the neutral rate for A, G polymorphisms in the genome (see previous paragraph). This result is incompatible with the hypotheses H2 and H3, as the frequency spectrum for the derived G-allele at non-coding edited sites should fit the neutral expectation (Table 2).

**Fig 1.**
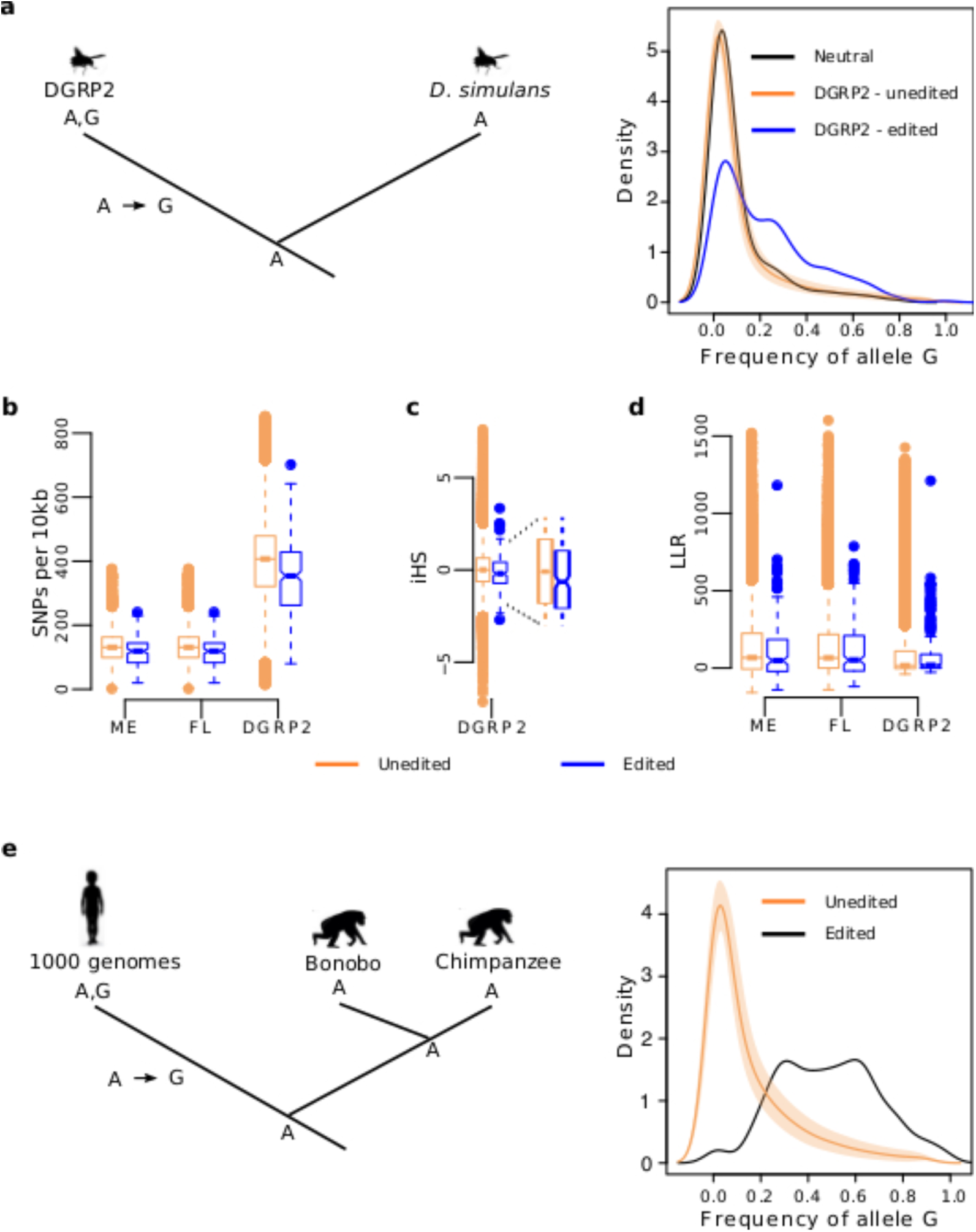
Properties of the G alleles segregating at edited sites in *D. melanogaster* and human. **a,** We used *D. simulans* as an outgroup to infer the ancestral state of the A,G polymorphisms in *D. melanogaster*. The right panel shows the average frequency spectrum and 95% confidence interval of the derived G alleles at unedited sites (peach) and the frequency spectrum for the derived G alleles at edited sites (blue). The shift of the blue distribution towards higher G allele frequencies is a signal of positive selection for the derived G alleles at edited sites. The black curve shows the expected frequency distribution of the derived G alleles at edited sites if they were neutral. **b,** Windows centered on polarized A-to-G mutations have lower diversity (in SNPs per 10kb) for edited SNPs than for unedited SNPs (*P* < 10^-4^ for each paired comparison; onesided Mann-Whitney-U test). **c,** At polarized edited sites, the extended homozygosity of the haplotype carrying the derived G allele is longer than that of the haplotypes carrying the ancestral A allele (average iHS score < 0). At unedited sites, the extended homozygosity is similar for both haplotypes (average iHS score ∼ 0). *P* = 0.004, one-sided Mann-Whitney-U test for the null hypothesis iHS (edited) ≥ iHS (unedited). **d,** The LLR comparing a long-term balancing selection model versus a neutral model tend to be lower for edited sites than for unedited sites (expected to be higher if balancing selection were more prominent for edited sites). *P* ≫ 0.05 for each paired comparison; two-sided Mann-Whitney-U test. **e,** We used Bonobo and Chimpanzee as an outgroup to infer the ancestral state of the genic A,G polymorphisms in the human genome. The right panel shows that G alleles segregate at higher frequencies in edited sites (black line) than in unedited sites (peach).

### Differentiated genomic footprints around edited and unedited sites in *Drosophila*

Two different scenarios may explain the higher frequency of the derived G allele at edited sites: directional selection in favor of the G allele or long-term balancing selection. We further looked for genomic signatures across the polarized polymorphisms that helped us to distinguish between these two scenarios.

According to the theory of selective sweeps, a new adaptive mutation appears on a single haplotype that quickly goes to fixation due to directional selection. The hallmark of a selective sweep is a reduction of nucleotide diversity near the adaptive mutation[22]. Accordingly, if the G allele at edited sites is positively selected, we expect reduced nucleotide diversity in genomic regions around polymorphic edited sites compared to unedited sites. We computed the number of single nucleotide polymorphisms (SNPs) in 10kb windows centered on edited A-to-G polymorphisms across the genome and tested whether these windows had the same nucleotide diversity than those centered on unedited A-to-G polymorphisms (Fig 1b). The average number of SNPs are 346, 125 and 116 for windows centered on edited sites (DGRP, FL and ME, respectively) and 398, 144 and 131 for windows centered on unedited sites (DGRP, FL and ME, respectively). Such a reduction of nucleotide diversity is significant in the three populations (*P* < 10^-4^ for each paired comparison; one-sided Mann-Whitney-U test) and a similar reduction of diversity is observed for 1kb windows (S3 Fig).

Another prediction of directional selection is that, because the adaptive G allele increases in frequency relatively fast, it will locate on an unusually long haplotype of low nucleotide diversity[23]. On the other hand, the haplotypes carrying the original A allele should be shorter than the haplotypes carrying the adaptive G allele but of similar length to haplotypes from a neutral genomic background. We used the genotypes of the 205 inbred lines from the DGRP2 to compute the integrated haplotype score (iHS)[23], an index that compares the extended homozygosity of the haplotypes carrying the derived G allele with that of the ancestral A allele. The iHS values at unedited A-to-G polymorphism (median iHS = 0.003) indicate that the haplotypes carrying the alleles at unedited SNPs have the same length and are likely neutral[23]. In contrast, the negative median iHS = −0.202 at edited A-to-G polymorphism (Fig 1c) indicate unusually long haplotypes carrying the derived G allele and suggest that these haplotypes have increased in frequency faster than neutral expectation. However, when testing one edited site at a time, only 12 of the iHS values are significant (*P* < 0.05, one-sided t-test for the null hypothesis iHS^edited^ ≤ iHS^unedited^), revealing the limitations of our analysis (see Discussion for further details).

The reduced nucleotide diversity near the edited A-to-G polymorphism and the longer haplotypes carrying the derived G alleles at edited sites is inconsistent with long term balancing selection, as a prediction of balancing selection is a local increase in nucleotide diversity[24]. To further evaluate long term balancing selection as one reason for the higher population frequency of the derived G allele at edited sites, we tested whether the local increase in nucleotide diversity relative to nucleotide divergence (i.e., fixed differences between species) is stronger near polymorphic edited sites than near polymorphic unedited sites[24]. To do so, we gathered a total of 100 nucleotide sites upstream and downstream of the polarized A-to-G polymorphisms across the genome, where a site is either a SNP or a fixed difference between *D. melanogaster* and *D. simulans*. For each window, we computed a log-likelihood ratio (LLR) that compares a balancing selection model against a neutral model based on the background genome pattern of polymorphisms[24]. Our analysis shows that the likelihood of the balancing selection model relative to that of the neutral model is lower in windows centered on A-to-G polymorphic edited sites than in windows centered on A-to-G polymorphic unedited sites (Fig 1d). The average LLRs comparing both models are 78, 120 and 111 for windows centered on A-to-G edited sites (DGRP2, FL and ME, respectively) and 83 and 136 for windows centered on A-to-G unedited sites (DGRP2 and both FL and ME, respectively). This result indicates that the signal of balancing selection is less prominent at A-to-G edited sites than at A-to-G unedited sites.

### Differentiated polymorphism pattern and allele frequency spectrum between edited and unedited sites of *Alu* repeats

We further applied our comparative analysis in humans to determine whether the selective footprints found in *Drosophila* were unique to this lineage or, otherwise common between these two distantly related species. Because the human genome is about two orders of magnitude larger than *Drosophila’s*, several difficulties arose, in particular: the list of (coding) edited sites is proportionally shorter than in *Drosophila* (in part due to the filtering by SNPs that is normally done to annotate the human editome) and the proportion of homologous nucleotide sites sequenced in other apes’ genomes (needed to polarize polymorphisms) is greatly reduced. Consequently, our approach in humans is inevitably more challenging and limited than in *Drosophila*. For instance, in our first attempt to apply our approach to humans, we integrated a recent list of 2,042 known coding edited sites[9] into a population genomics database compiled from the 1,000 Genomes Project[25] and the Great Ape Genome Project[26]. However, only 10 of the 2,042 edited sites were represented in our database, impeding any further genome-wide analysis.

Because humans have more than a million copies of *Alu*[27] and virtually all adenosines within *Alu* repeats that form double-stranded RNA undergo A-to-I editing[28], we used our population genomic approach on *Alus*. By using *Alus* we are limiting our analysis to silent (most genic *Alu* repeats occur in introns and 3’ UTRs) and intergenic A sites, but we gain in numbers enough to look for genome-wide polymorphism patterns. With this in mind, we analyzed RNA-Seq data from 105 control (healthy) breast samples from The Cancer Genome Atlas (TCGA) and annotated *de novo* a list of 28,322 highly-edited sites at *Alu* repeats, 1,838 of them represented in our database (1,208 genic and 630 intergenic; Table 2). Remarkably, we found a 3-fold increase in the nucleotide polymorphism at edited *Alu* sites (19%) compared to unedited *Alu* A-sites (6%) located in genes. In addition, the G nucleotide is the alternative allele in 97% of the polymorphic edited sites, but only in 58% of the unedited polymorphic sites (Table 2). We used chimpanzee and bonobo population genomic data to infer the ancestral state of the A,G polymorphisms occurring at genic *Alus*, and compared the frequency spectrum of the derived G alleles segregating at edited and unedited sites. Fig 1e shows that derived G alleles at edited sites segregate at higher frequency than derived G alleles at unedited sites. Notably, we observed a similar nucleotide polymorphism pattern (Table 2) and allele frequency spectrum (S5 Fig) for edited sites in intergenic *Alu* repeats. Our study in humans therefore confirms our results in *Drosophila* and suggest that a significant fraction of A-to-G mutations at edited sites is also adaptive in humans, including those occurring in intergenic regions.

## Discussion

The binary classification (edited/unedited) of *Drosophila* and human population genomic data based on a posttranscriptional modification uncovered an evolutionary footprint that, otherwise, would remain hidden. Several of these footprints seem incompatible with the current hypotheses on the evolution of A-to-I RNA editing and prompt us to suggest an additional hypothesis that may better explain our results.

The extraordinary differences of the polymorphic rates and polymorphism types between edited and unedited sites are very unlikely affected by differences in the usage of synonymous codons (Fig 2a), gene expression level (Fig 2b) or recombination rates (Fig 2c and S4 Fig) between edited and unedited sites. Higher GC biased gene conversion (i.e., the unequal exchange of genetic material between homologous loci) is also an unlikely source of bias as there is no GC biased gene conversion in *Drosophila*[29] and we restricted our analysis in human to A-sites of *Alu* elements, ensuring identical local sequence for both edited and unedited sites. In addition, we did no find significant differences in the nucleotide composition around edited and unedited A-sites in *D. melanogaster* that might suggest context-driven local mutation rates (Fig 2d). Finally, we found similar results for *Drosophila* and human out of different editing annotation strategies and population genomic datasets, suggesting that annotation artifacts are not likely affecting our analysis.

**Fig 2.**
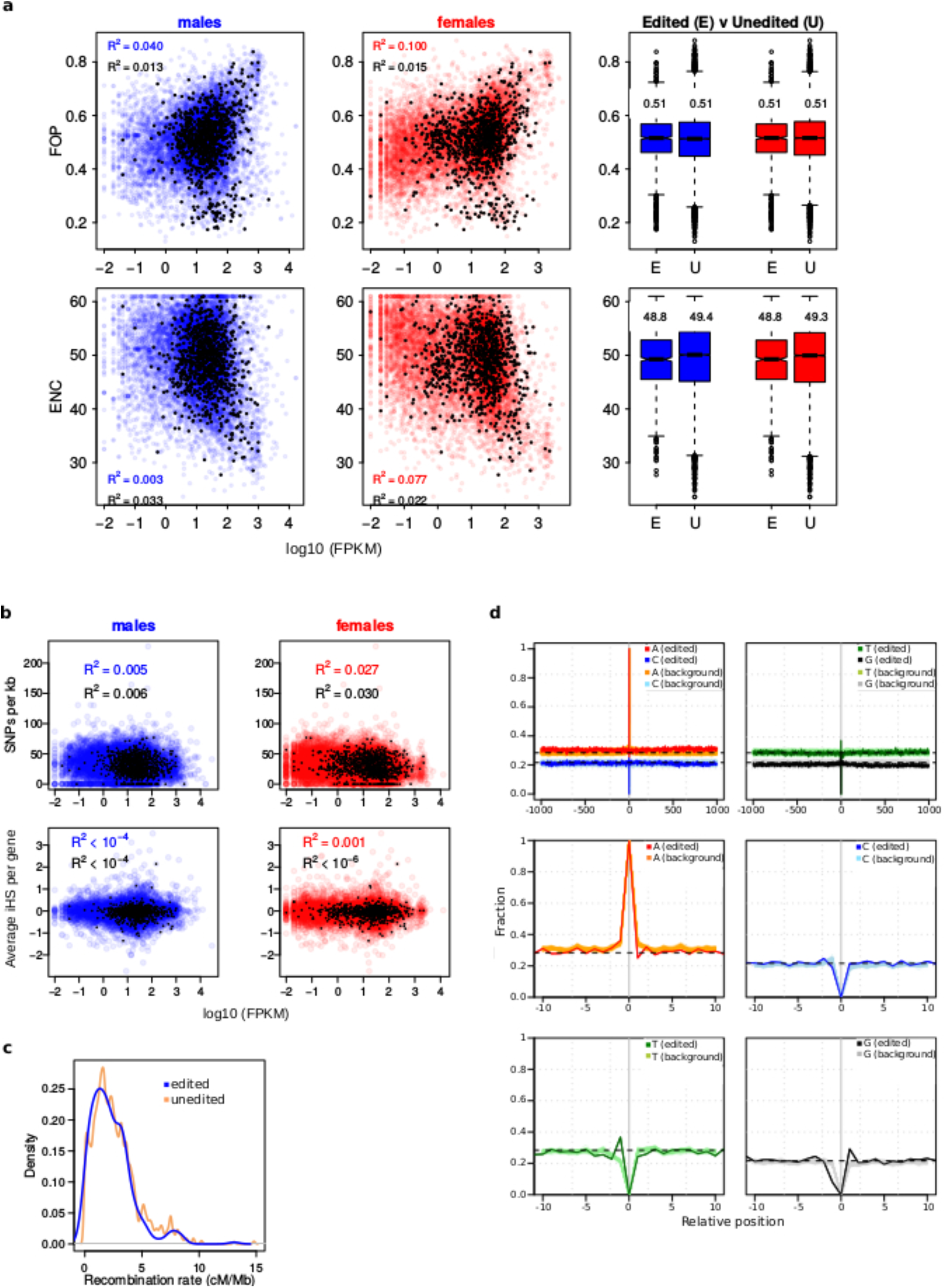
Control analyses for differences in polymorphic rates and polymorphism types as a byproduct of gene expression level, recombination rate and local sequence composition in *Drosophila*. **a,** Bias in synonymous codon usage per gene is represented as a function of gene expression level in males (blue) and females (red). Gene expression level only explains 4% (males) to 10% (females) of the total variance in codon bias when measured as the frequency of optimal codons (FOP; the higher, the more biased) and 0.3% (males) to 7% (females) of the total variance in codon bias when measured as the effective number of codons (ENC; the lower, the more biased). The coefficient of determination for edited sites (black dots) is even lower than for unedited sites. Numbers in the boxplots refer to the mean. **b,** Nucleotide diversity (SNPs per kb per gene) and iHS (averaged per gene) does not correlate with gene expression level. Black dots: genes containing edited sites. Blue and red dots: unedited genes. **c,** Local recombination rates in 10 kb windows centered on edited (blue) and on unedited (peach) sites show identical distributions. **d,** Nucleotide profiles show that local sequence context around edited and unedited sites (±1000 bp and ±10 bp) are virtually identical.

The fact that the nucleotides C and T are virtually absent at edited sites suggest strong functional constraints upon edited A-sites in humans and flies. This implies that the relative fitness (s) of edited A-sites is much higher than that of the alternative C and T alleles (*s_A_* ≫ *s_C,T_*). In addition, the fact that derived G alleles at edited A-sites segregate at higher frequencies than expected (Fig 1a and 1e) indicates that the A-to-G mutations at edited sites are generally adaptive. In other words: *s_G_* > *s_A_* ≫ *s_C,T_* at edited sites. These two observations are also difficult to explain according to the current hypotheses on editing and shed light on the adaptive roles of the G mutations at edited sites and on the A-to-I RNA editing itself. Our hypothesis is that a genomically encoded G nucleotide is generally adaptive at edited sites because it mimics the function of the edited RNA. This implies that A-to-I RNA editing is also generally adaptive (hypothesis H4, Table 1). If A-to-I RNA editing were not adaptive, the G allele would not reveal signatures of adaptation and C and T alleles would be also found at edited SNPs (both coding and non-coding).

We showed that directional selection in favor of the derived G allele is more likely than balancing selection acting at A,G polymorphic edited sites. However, the evidence is weak for several reasons. First, we can only analyze incomplete selective sweeps because we do not know which G nucleotide sites currently fixed in *D. melanogaster* were edited A-sites in the past. Second, the selection strength may depend on the dominance of the derived G allele. For instance, it is likely that the dominance has a more prominent effect at nonsynonymous G mutations than at silent mutations. Third, although directional selection may be more prominent, balancing selection may still occur at some edited sites. Despite these limitations, by averaging over many sites, the footprint for directional selection, and not balancing selection, becomes more evident (but not conclusive).

The adaptive potential of A-to-I RNA editing by modifying the protein sequence have been recently proven. Garrett and Rosenthal[30] showed that the editing level of the mRNA encoding the octopus’ potassium Kv1 channels correlates with the water temperature where the octopus’ species were captured. Most importantly, a concomitant physiological amelioration at cold Antarctic temperatures indicates that RNA editing may play a significant role in thermal adaptation in this species. The important role of A-to-I RNA editing on posttranscriptional regulation, including editing of genic *Alu* sequences[1], also suggest an adaptive potential of editing as a checkpoint to gene expression control. In summary, the adaptive role of the G mutation at edited sites may come in two ways: by encoding the same protein variant and “encoding” the same RNA secondary structure as in the edited RNA.

The adaptive role of the G mutations at edited A-sites of intergenic *Alu* repeats is less obvious to explain. It has been shown that ADAR1 mutants over-express genes containing edited *Alu* repeats and that *Alu* editing is involved in the nuclear retention of the cognate mRNA[31]. We suggest that A-to-I RNA editing (and A-to-G mutations mimicking the editing function) might be an adaptive mechanism to prevent the deleterious effect of retrotransposition of intergenic *Alu* repeats and could work in two flavors: 1) by silencing the expression of the *Alu* repeats or 2) by retaining the transcribed *Alu* repeats to impede their retrotranscription in the cytoplasm.

We expect that new population genomics data and new editome annotations will help us to find additional signs of positive selection in other animal classes and confirm the pervasive adaptive potential that A-to-I RNA editing offers to these two distantly related species, *D. melanogaster* and human. Our novel approach will hopefully help to expose similar genome-wide adaptive patterns associated with the expanding epitranscriptome landscape.

## Methods

### Population genomic data

We downloaded the genotypes of the 205 inbred lines annotated in the *Drosophila* Genetic Reference Panel 2[13] (http://dgrp2.gnets.ncsu.edu/). In addition, we also analyzed pooled DNA-Seq data from *D. melanogaster* flies collected in 2010 from outbred populations in Maine (86 lines) and Florida (39 lines)[14]. We trimmed 101 bp paired-end reads with ConDeTri[32] using the following parameters: hq=20, lq=10, frac=0.8, minlen=50, mh=5, ml=1, and mapped with NextGenMap[33] the remaining reads longer than 50 bp to the *D. melanogaster* reference genome, release r5.40 (ftp://ftp.flybase.net/genomes/). Next, we removed reads with a mapping quality value lower than 20 with SAMtools[34]. We called SNPs for each dataset when the coverage was ≥ 10 at this nucleotide site and at least two reads carried the alternative allele.

A pileup from 6 *D. simulans’* sequenced genomes was downloaded from the *Drosophila* Population Genomics Project (http://www.dpgp.org/). We used UCSC’s liftover tool[35] to convert dm2 coordinates into dm3 coordinates (BDGP Release 5).

Primate population genomic data was downloaded from the Great Ape Genome Project[26]. We converted the coordinates from hg18 to hg19 using liftover and used hg19 nucleotide site ID to merge the Great Ape population genomics data with the human data from the 1,000 Genomes Project[25]. The merged population genomics database consists of 179,546,112 entries indicating homologous nucleotide sites in great apes and allele frequency information in humans.

### A-to-I RNA editing data

We used the latest annotation of the A-to-I RNA editing sites in *D. melanogaster*, which consists of 3,581 sites[8]. In this study, editing events were called when G allele expression was detected from a homozygous AA genotype. The potential editing sites were further confirmed by the absence of G allele expression at putative editing sites in ADAR^−/−^ mutants generated from the same isogenic line.

We annotated *de novo* the A-to-I RNA editing sites occurring in *Alu* repeats in a conservative way. Briefly, we mapped RNA-Seq data from 105 control (healthy) breast tissue samples available at The Cancer Genome Atlas (TCGA) project (http://cancergenome.nih.gov/) against the human reference genome (hg19) with STAR aligner v2.3.0[36]. Only uniquely mapped reads with less than 5% mismatches were kept for further analysis, allowing us to test a total of 148,961,882 A sites for A-to-I RNA editing. For the purpose of this study, we defined a site to be edited if 1) the G allele were found at >1% of the reads in >50% of the breast samples and 2) the G allele was not found in the dbSNP (build 146) at frequency >0.5. Otherwise, the A site was defined as unedited. This definition allowed us to detect 28,322 highly edited sites out of the ∼149 million A sites tested.

### Polarizing A-to-G mutations in D. melanogaster and human

We downloaded pairwise *D. melanogaster/D.simulans* axt alignment files from UCSC (http://hgdownload.soe.ucsc.edu/goldenPath/dm3/vsDroSim1/). A script was generated to parse the alignment files and detect the homologous sites in *D. simulans* reference genome and in six additional *D. simulans* genomes downloaded from the *Drosophila* Population Genomics Project (http://www.dpgp.org/). A-to-G mutations were inferred to occur on the *D. melanogaster* lineage (DGRP, ME and FL populations) when the homologous site in the *D. simulans* lines was A (i.e., monomorphic in *D. simulans* population).

We parsed the pileup file from the Great Ape Genome Project and compiled the list of human A,G SNPs that likely originated by A-to-G mutation in the human lineage. The ancestral state of an A,G polymorphism was already inferred in the original study and stored in the pileup file as *node* 18[26].

### Allele frequency spectrum

Low coverage in pool-sequencing experiments may inflate the frequency estimation of alleles segregating at low frequencies. We tested for different coverage among edited and unedited polymorphisms and for a correlation between coverage and minor allele frequency in ME and FL populations. S2 Fig shows that the coverage is not different between edited and unedited sites and that allele frequency and coverage do not correlate. Therefore, we are confident that the higher frequency of the G allele in edited sites is not due to an artifact associated with coverage.

After polarizing the polymorphism data with *D. simulans*, we found 462,801, 110,844 and 125,807 A-to-G polymorphic sites in DGRP2, ME and FL populations, respectively, that most likely originated from A-to-G mutations. 303, 155 and 179 of these sites are edited sites in DGRP2, ME and FL populations, respectively (S2 Table).

For DGRP2 data, we computed the frequency of the derived G allele as 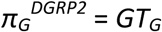 / (*GT_G_ + GT_A_*), were *GT_G_* and *GT_A_* are the number of lines with genotype GG and genotype AA, respectively. For ME and FL populations, we computed the frequency of the G allele as 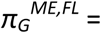 *g* / *r*, as suggested for pool-sequencing data[37], where *g* is the number of DNA-Seq reads carrying the G allele and *r* is the total number of reads mapped at this site. To compute the allele frequency spectrum of the derived G alleles across the genome, we sampled 303, 155 and 179 sites from the 462,801, 110,844 and 125,807 polarized A-to-G polymorphic sites in DGRP2, ME and FL populations, respectively. We repeated the sampling 100,000 times (per population) to compute the average distribution and the 95% confidence interval for each frequency class. The expected neutral allele frequency spectrum of the G alleles segregating at the edited sites was computed by plugging the 303, 155 and 179 allele frequencies into Kimura and Crow’s formula[38]

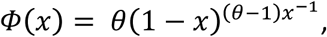

where *x* is the allele frequency and *θ* = 4*N_e_v*. We used *θ* = 0.007, as previously estimated for DGRP2[13,39], and ME and FL populations[14]. The expected neutral allele frequency spectrum fits the observed frequency spectrum of the 462,801, 110,844 and 125,807 polarized unedited sites in DGRP2 (Fig 1a), ME and FL populations (S1 Fig). To plot the neutral allele frequency spectrum for Fig 1a, we only considered G alleles segregating at frequencies higher that 1% and lower than 99%.

We polarized 176,311 tested A,G human polymorphisms occurring at genes that most likely originated from A-to-G mutations; 231 of them corresponded to edited sites in genes (Table 2). To compute the allele frequency spectrum of the G allele at genes, we sampled 231 sites from the 176,311 unedited A,G polymorphisms. We repeated the sampling 100,000 times and compute the average allele frequency spectrum and the 95% confidence interval for each frequency class. We took the frequency of the G alleles from the 1,000 Genomes Project. With regards to intergenic regions, we polarized 196,140 tested A,G human polymorphisms that most likely originated from A-to-G mutations; 110 of them corresponded to edited sites (Table 2). The sampling procedure was as explained for genic A,G polymorphism with sampling size 110.

### Testing for balancing selection and directional selection

To test for directional selection in favor of the derived G allele in edited sites, we first tested whether diversity was lower around edited sites than around unedited sites. To this aim, we counted the number of SNPs in windows of 10kb centered on each polarized A-to-G polymorphism. The ancestral allele was again determined based on data from *D. simulans*. We also used the recombination rate data from Ref.[40] to linearly interpolate local recombination for the 10kb windows. The distribution of local recombination rates at edited and unedited sites are essentially identical (S4 Fig), ruling out a bias in our diversity analyses caused by differences in recombination rates between edited and unedited sites.

We also computed the integrated haplotype score (iHS)[23] using the software rehh[41] as a second approach to test for directional selection in favor of the derived G allele in edited sites. G alleles raising rapidly due to strong selection will have less chances to accumulate new mutations around and will tend to have high levels of haplotype homozygosity extending much further than expected under a neutral model. The rationale of the iHS approach is therefore to test whether the derived G allele at an edited site tends to segregate on an unusually long haplotype of low diversity[23]. Because haplotypes cannot be inferred for pool-sequencing, we computed iHS only for the DGRP2 population. Negative values of iHS indicate unusually long haplotypes carrying the derived G allele compared to the ancestral A allele. Values of iHS close to zero indicate that the haplotypes carrying both the ancestral and the derived alleles are equally large and the tested SNP is likely neutral[23].

To scan for polymorphic sites under balancing selection, we used the software ballet[24]. Ballet combines intraspecies polymorphism and interspecies divergence with the spatial distribution of polymorphisms and substitutions around a selected site. The signature of balancing selection is that of a local increase in diversity relative to divergence, and a skew of the site frequency spectrum towards intermediate frequencies. The method outperforms both the HKA test and Tajima’s D under a diverse set of demographic assumptions, such as a population bottleneck and growth[24]. We calculated a log-likelihood ratio (LLR) for each polymorphic site implemented in the test type T1. The input files for ME and FL population consisted of the polymorphic state inferred from the pool-sequencing data. Because ballet can only handle a maximum of 100 lines, we used a random sample of 50 isogenic DGRP2 lines (Fig 1d) and of 100 randomly sampled lines to carry out the LLR computation. The result obtained for 100 lines are similar to the result for 50 lines (not shown). We specified a window size of 200 sites, as little is gained by incorporating information from additional sites[24], where a site is an intraspecies polymorphism or a divergent site. Divergent sites to *D. simulans* were defined as single nucleotide substitution: i.e., homologous non-polymorphic (fixed) sites that contain different nucleotides between *D. melanogaster* and *D. simulans*. Ballet also utilizes information regarding the recombination distance between sites. We used the recombination rate data from Ref.[40] to linearly interpolate recombination distance between two consecutive sites.

### Estimation of f_s_ and f_n_

To estimate *f_s_* and *f_n_* in *D. melanogaster*, we first compiled all A sites from the reference genome, release r5.40, and generated a variant call file with all potential A,G polymorphisms. We used this file as input to CooVar[42], which analyzed the effect of each A-to-G mutation in coding regions. The output files were integrated into the DGRP2, FL and ME polymorphism database to identify the potential A,G synonymous and nonsynonymous polymorphism that are actual A,G polymorphisms.

### Gene expression and codon usage data

We download gene expression data from the GEO (acc. GSE67505). The expression data was obtained from pooled RNA-Seq data for the DGRP2 lines, as described in the original study[43]. The published expression tables are given separately for male and females in FPKM units. To test for correlation between gene expression levels and non-random usage of codons (i.e., codon bias), we downloaded two measurements of codon bias (the effective number of codons or ENC and the frequency of optimal codons or FOP) from the sebida database[44] and fused the DGRP2 expression data with sebida data by means of the FlyBase gene IDs. Genes containing at least one edited site were coined edited genes and unedited genes otherwise.

### Nucleotide profiles

The nucleotide profile around edited sites was calculated as the fraction of A, C, G and T nucleotides at each nucleotide site upstream and downstream (±10 bp and ±1,000 bp) the edited site. For the background data, we sampled *a* = 1,657 genic A sites and *t* = 1,549 T sites from the *D. melanogaster* genome, where *a* and *t* are the number of annotated edited sites in the direct and inverted strands, respectively, and repeated this operation 100 times to compute the fraction of each nucleotide type at each nucleotide position upstream and downstream the sampled A/T unedited sites.

## Data availability

Computer code and data is available upon request to the authors.

## Acknowledgments

We thank Angela M. Hancock and Michael DeGiorgio for valuable comments and suggestions, Nadia Singh for providing data and Jennifer Gage for proofreading the manuscript. Funded by Austrian Science Foundation grant SFB F43-13 and P26882 (M.J.) and Israel Science Foundation grant 379/12 (E.E.). Author contributions: MJ, AvH and MG conceived the project. MG designed the experiments. NP, CDH, I.B, E.E and MG analyzed the data. MG wrote the paper. NP, CDH, I.B, E.E, MJ, AvH and MG revised the paper.

